# REMI-seq: Development of methods and resources for functional genomics in *Dictyostelium*

**DOI:** 10.1101/582072

**Authors:** Nicole Gruenheit, Amy Baldwin, Balint Stewart, Sarah Jaques, Thomas Keller, Katie Parkinson, Rex Chisholm, Adrian Harwood, Christopher R. L. Thompson

## Abstract

Genomes can be sequenced with relative ease, but ascribing gene function remains a major challenge. Genetically tractable model systems are crucial to meet this challenge. One powerful model is the social amoeba *Dictyostelium discoideum*, a eukaryotic microbe widely used to study diverse questions in cell, developmental and evolutionary biology. However, its utility is hampered by the inefficiency with which sequence, transcriptome or proteome variation can be linked to phenotype. To address this, we have developed methods (REMI-seq) to (1) generate a near genome-wide resource of individual mutants (2) allow large-scale parallel phenotyping. We demonstrate that integrating these resources allows novel regulators of cell migration, phagocytosis and macropinocytosis to be rapidly identified. Therefore, these methods and resources provide a step change for high throughput gene discovery in a key model system, and the study of genes affecting traits associated with higher eukaryotes.

## Introduction

The past two decades have seen an exponential increase in genome sequence availability. However, our ability to comprehensively ascribe function to gene sequences is still limited. One solution is to identify and characterise the effects of loss of function mutations in all genes across the whole genome. Consequently, efficient tools for random or systematic genome wide mutagenesis are crucial and are now readily available in many model organisms. When coupled with methods that allow high throughput characterisation of these mutants, this has the potential to provide systems level genotype-phenotype information.

Microbial systems have proven particularly powerful models for such genome-wide functional analysis. First, they tend to have small and mostly haploid genomes, which allows libraries of thousands of ‘barcoded’ single-gene loss or gain of-function mutants to be generated by homologous recombination (Giaever *et al.*, 2002) or large scale random insertional mutagenesis (Hayes, 2003; van Opijnen, Bodi and Camilli, 2009; Akerley *et al.*, 1998; Zhang *et al.*, 2014). Second, their fast generation times and relatively simple phenotypes allow genome-wide forward genetics approaches in which these libraries of mutants can be screened under different selection regimes to enrich for advantageous mutations or deplete disadvantageous mutations. In recent years, massively parallel sequencing has allowed the frequency of every mutant allele to be quantified in a library before and after a selection (van Opijnen, Lazinski and Camilli, 2015). For example, this approach has been applied to define genes required for growth, survival under different conditions, virulence and antibiotic resistance, as well as for genetic interaction discovery (van Opijnen, Bodi and Camilli, 2009; Zhang *et al.*, 2012; Goodman *et al.*, 2009; Gawronski *et al.*, 2009; Mann *et al.*, 2012; Gallagher, Shendure and Manoil, 2011).

Functional genomic studies in simple microbial systems illustrate the power of parallel phenotype analyses for understanding biological systems. However, one drawback is that because most microbial systems are unicellular, they are often limited by the spectrum of phenotypes that can be studied. In addition, comprehensive phenotypic analyses of multicellular traits characteristic of higher eukaryotes, such as cell differentiation, cell-cell signalling and cell migration are challenging. First, large diploid genomes provide an obstacle to generating extensive mutant collections. Large numbers of individual mutant lines must be generated and then bred to homozygosity, or alternatively strains that express inhibitors or activators of gene function must be generated. Second, phenotypic analyses are experimentally challenging, and individual strains often must be grown up and examined. Whilst cell culture mutants have been generated in some model systems, this in turn limits the spectrum of phenotypes that can be studied. Despite these limitations, the availability of mutant collections, together with sophisticated screening methods has undoubtedly underpinned our ability to use model systems to better understand gene function in diverse biological processes.

*D. discoideum* is a well-established eukaryotic microbial model for studying diverse processes in cell and developmental biology because it exhibits a unique lifecycle, with both unicellular and multicellular stages (Kessin, 2001). During unicellular growth, it is motile, actively seeking and engulfing bacteria via actin-mediated locomotion and phagocytosis. It divides by binary fission, again using an actin mediated process, and can grow in liquid medium by up-regulation of macropinocytosis. Upon starvation, it initiates a programme of multicellular development, where approximately 100,000 cells aggregate by chemotaxis to cAMP. As it enters its multicellular state, cell-cell signalling controls the temporal and spatial patterning of a small number of different cell types. Ultimately, these cells terminally differentiate to form a fruiting body, composed of dead stalk cells that aid the dispersal of viable spores. As a consequence, the *Dictyostelium* life cycle provides a powerful model for studying fundamental cell processes, such as cell motility, chemotaxis (Nichols, Veltman and Kay, 2015), macropinocytosis, and phagocytosis (King *et al.*, 2013; Cornillon *et al.*, 2000); cell-cell signalling, differentiation, morphogenesis during multicellular development; and longer term evolutionary changes due to social conflict (Gruenheit *et al.*, 2017; Strassmann and Queller, 2011; Kuzdzal-Fick, Queller and Strassmann, 2010; Foster *et al.*, 2004; Wolf *et al.*, 2015; Madgwick *et al.*, 2018). Unsurprisingly, sequencing of the compact haploid (34Mbp) genome reveals widespread gene conservation with higher eukaryotes (Eichinger *et al.*, 2005). Finally, random mutagenesis by restriction enzyme mediated integration (REMI) insertion (Kuspa and Loomis, 1992) has been developed to allow forward genetic identification of novel components of different biological processes (Santorelli *et al.*, 2008; Khare *et al.*, 2009; Parkinson *et al.*, 2011; Kibler *et al.*, 2003; Chattwood *et al.*, 2013; Thompson *et al.*, 2004; Adachi *et al.*, 1994; Thomason, King and Insall, 2017).

Many studies illustrate the potential of *Dictyostelium* as a model for understanding conserved biological processes. However, its true power for functional genomics has yet to be harnessed. Whilst genomic, proteomic and transcriptomic studies can be performed with relative ease, testing the functional importance of identified genes is challenging due to a lack of a genome wide collection of mutants. Individual mutants can be made by gene targeting via homologous recombination (De Lozanne and Spudich, 1987; Paschke *et al.*, 2018; Katz and Ratner, 1988) or CRISPR/Cas9 technology (Sekine, Kawata and Muramoto, 2018). However, such gene-by-gene approaches are time consuming and resource intensive. Large-scale generation of REMI mutants has always been possible, but has been limited by the need to identify individual insertion sites after genetic screens (Keim, Williams and Harwood, 2004; Adachi *et al.*, 1994).

To address these problems, we have developed a method, REMI-seq, for high throughput parallel identification of REMI insertion points. This allowed us to generate a near genome wide collection of individual REMI mutants, providing a new resource for the research community. This resource can be utilised in two modalities. First, the Genome Wide *Dictyostelium* Insertion (GWDI) bank is a gridded-library of 21,529 different mutants, with each mutant catalogued and available for research analysis as individual or groups of mutants, thus providing a link between existing ‘omics based’ resources. Second, we have generated a large pool of sequenced ‘barcoded’ mutants that enables large-scale parallel phenotyping. As a proof of principle, we performed parallel selections for growth on bacteria and liquid medium, where phagocytosis and macropinocytosis respectively are used as the major routes of nutrient uptake. Dynamic changes in relative abundance of each mutant allele within the population were measured by deep sequencing to identify advantageous and disadvantageous mutants under different conditions. These data reveal that the use of functional outcomes reveals novel biological pathways that mediate cell behaviour. Together, the development of these resources provide an important advance as they will aid the high-throughput forward and reverse genetic identification of genes affecting traits associated with higher eukaryotes.

## Results and discussion

### REMI-seq: A method for genome-wide gene disruption and mutant identification

We adapted the standard REMI protocol to create a methodology for genome-wide mutagenesis that we termed REMI-seq. This allows high-throughput insertional mutagenesis and rapid mutant identification. For mutagenesis, we designed a series of plasmids (pGWDI) (Figure 1A, Supplementary File 1), from which PCR was used to generate integration DNA fragments containing a blasticidin resistance cassette (Figure 1B, Supplementary File 2). These fragments were introduced into the *D. discoideum* laboratory strain AX4 together with the appropriate accompanying restriction enzyme. Different versions of the pGWDI fragments can target either DpnII (GATC site; pGWDI-G) or NlaIII (CATG site; pGWDI-C) sites. These restriction sites were chosen because they are overrepresented in intragenic sequences. 58,852 out of 69,422 (85%) of DpnII sites and 41,845 out of 49,365 (85%) of NlaIII sites occur within coding regions of genes (Supplementary File 3). From a total of 13,412 putative genes, 12,391 genes contain one or more DpnII or NlaIII sites (or both) (Supplementary File 3). Consequently, more than 92% of the annotated genes can be targeted using a combination of the two insertion vectors. Furthermore, of the remaining 1,021 genes containing neither site, 496 contain restriction sites in their respective promoter regions (< 500 bp upstream of the start codon). Of the remaining 525 genes, 116 are tRNAs, 28 are small non-coding RNAs, and another 35 are pseudogenes or transposable elements. Finally, these sites are uniformly distributed across the genome (Figure 1C), with mean numbers of 202 GATC and 143 CATG sites per 100,000 bp interval. Only four and three intervals show a significantly higher number of DpnII or NlaII sites respectively (3 standard deviations higher than the mean), and thus represent putative hotspots.

**Figure 1.**
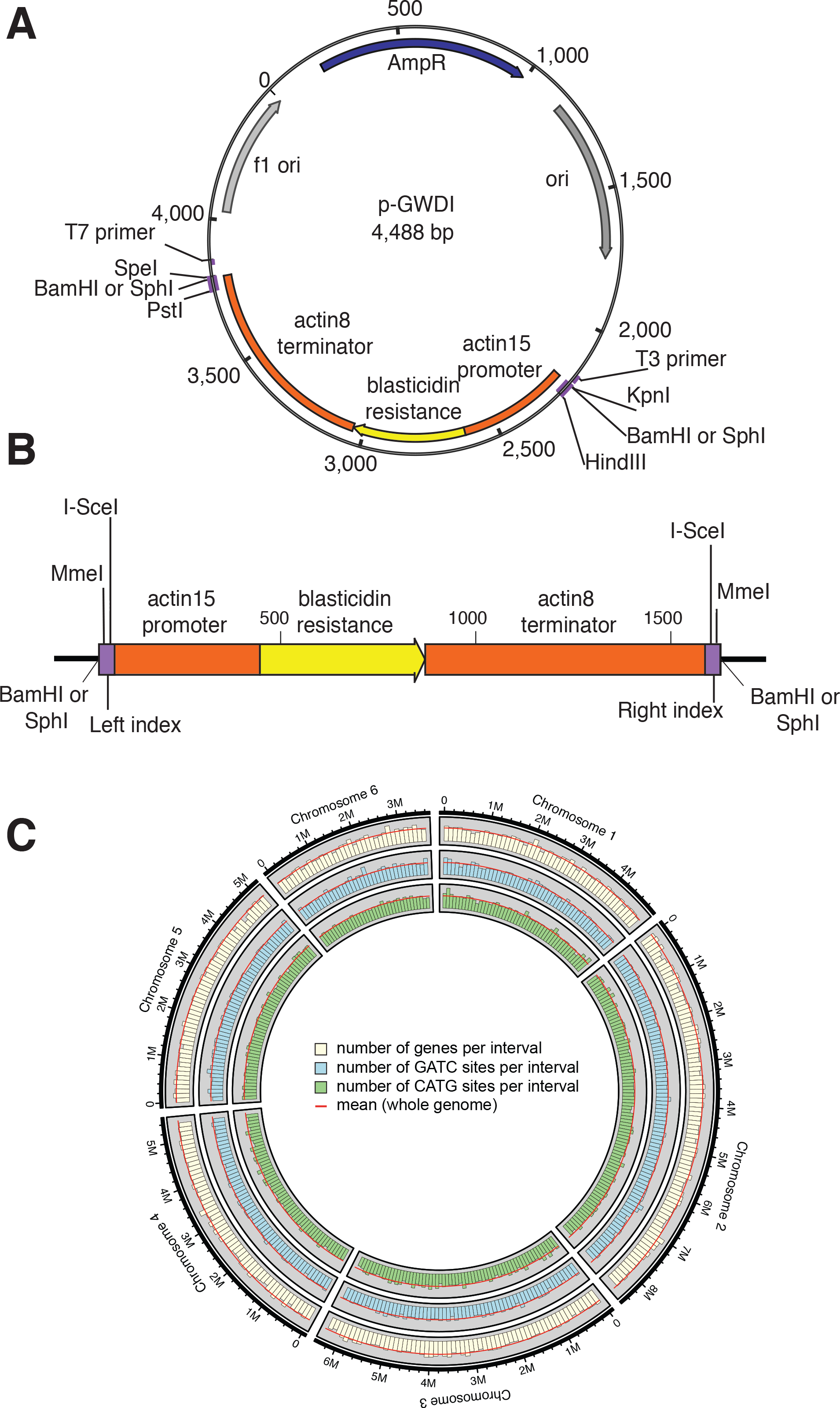
A modified REMI vector for genome wide gene disruption in *Dictyostelium*. **A. Schematic of the REMI-seq vector.** pGWDI is a derivative of pLPBLP which encodes the blasticidin S deaminase selectable marker under the control of the actin15 promoter and actin8 terminator sequences. The sequence was modified (purple box) to contain sequences for REMI insertion, identification of insertion point. The pGWDI-C and pGWDI-G plasmids are identical with the exception that they have SphI and BamHI sites for integration. In addition, four different versions of each vector were generated with a unique index sequence. **B. Schematic of amplified Insert for REMI mutagenesis**. The insert was generated by the amplification of the pGWDI template between the T7 and T3 primer sequences. The PCR product was cleaved with BamHI or SphI to generate sticky ends. The I-SceI and MmeI sites are required to capture the insert - genomic DNA junction for analysis via next generation sequencing. The left and right indices allow increased multiplexing for insertion point identification. **C. Computational predictions suggest REMI mutagenesis is random.** Predicted SphI and BamHI sites are randomly distributed across the genome. The density of each site was mapped onto 10kb fragments of the genome. Few regions are under- or overrepresented, and thus represent putative cold or hotspots.

To identify the insertion sites in an efficient and inexpensive fashion, we developed a novel strategy (REMI-seq) (Figure 2), based on a similar strategy used by Tn-seq (van Opijnen, Bodi and Camilli, 2009). The termini of the GWDI insertion fragment possess recognition sequences for the type IIS restriction enzyme MmeI, which cuts 19-20 bp downstream of the recognition site. A cut site for the meganuclease I-SceI was positioned 3’ to each MmeI site, which does not appear in the *D. discoideum* genome, (Figure 1B and Figure 2). Consequently, when genomic DNA is digested with MmeI, it will cut within the genomic DNA flanking the site of insertion (Figure 2), and subsequent digestion of genomic DNA with I-SceI will result in the generation of two 47 bp fragments. These fragments can be enriched by PCR, purified and sequenced by Illumina sequencing (Figure 2), with each read containing a 28 bp sequence comprised of the I-SceI and the MmeI restriction sites, vector identifier index sequence, the remnants of the DpnII or NlaIII site, and importantly an additional 19-20 bp of genomic sequence derived from the site of insertion. Even though the tags are fairly small, in silico analysis of the *Dictyostelium* genome revealed that most genes (11,244 out of 13,421) have at least one uniquely identifiable tag (Supplementary File 3). Consequently, the REMI-seq method has the potential to allow the high throughput generation of a near genome wide insertion mutant resource in *Dictyostelium*.

**Figure 2.**
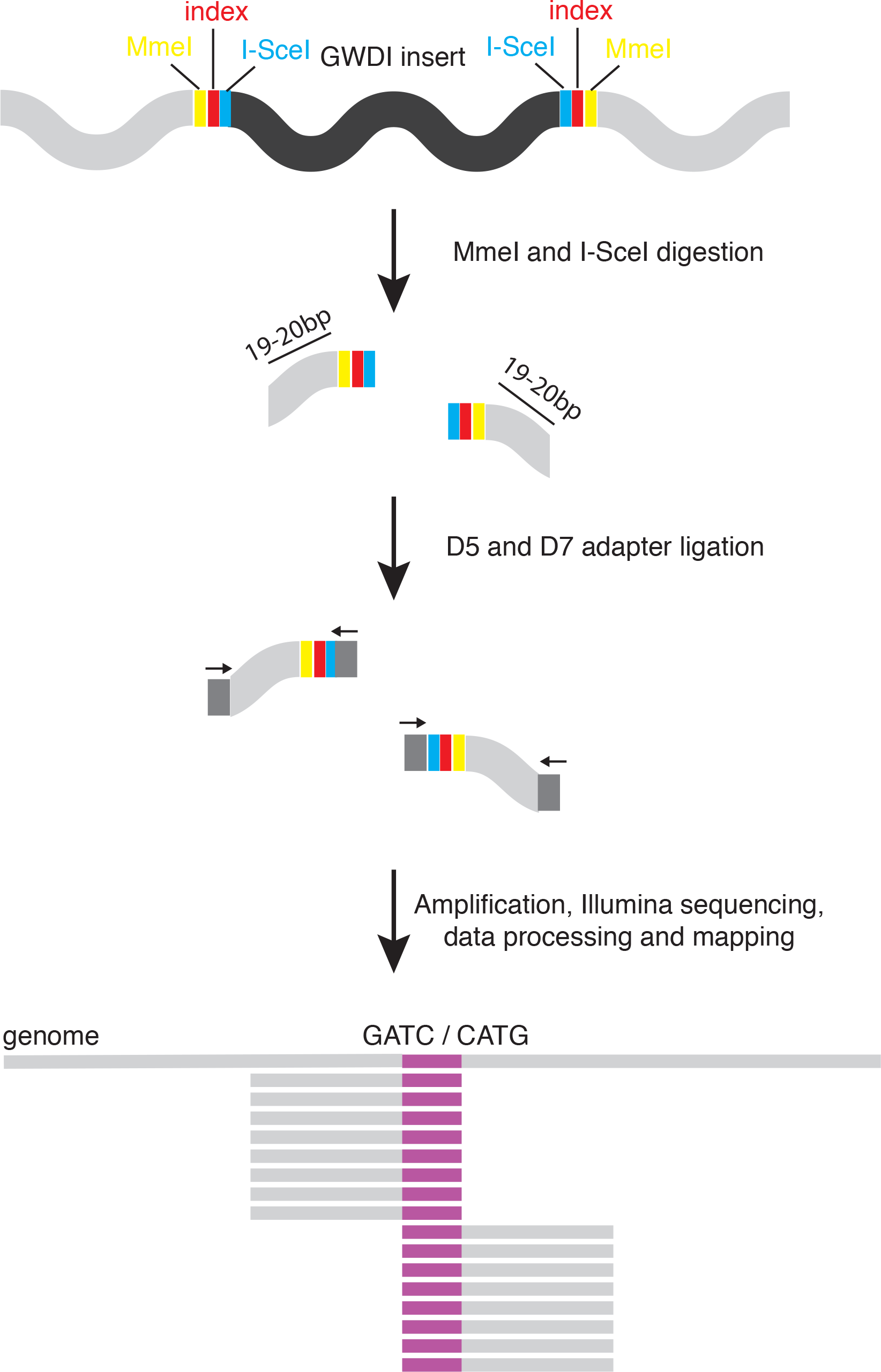
The REMI-seq method for insertion point identification. The GWDI fragment containing a blasticidin resistance cassette (black) has inserted into the *Dictyostelium* genome. The termini of the GWDI insertion fragment were engineered to contain recognition sequences for the type II restriction enzyme MmeI (TCCRAC) (yellow), and two sites for the meganuclease I-SceI (TAGGGATAACAGGGTAAT) (blue), which does not appear in the *Dictyostelium* genome. If gDNA is digested with MmeI (which cuts 19-20 bp downstream of the recognition site and leaves a two base pair 3’ NN overhang) and I-SceI (which leaves a four base pair 3’ ATAA), this results in the generation of two 47-48bp fragments. These fragments can be enriched and separated from other MmeI genomic fragments by PCR and gel extraction, thus allowing identification of the insertion site by sequencing. Therefore, sequencing reads contain 28 bp of sequence comprised of the I-SceI and the MmeI restriction sites, vector tag (red), remnants of the genomic DpnII/NlaIII site, as well as 19/20 bp of genomic sequence at the site of insertion.

### GWDI-bank: Isolation and identification of large numbers of *Dictyostelium* REMI mutants

To date, the number of available mutants (n = 723) in *Dictyostelium*, represents only 6% of the total genome. Furthermore, these mutants were made in a variety of different genetic backgrounds, making comparisons difficult. We thus applied REMI-seq to generate thousands of new mutants in the same isogenic background. In our pipeline (Figure 3), multiple independent transformations were first performed resulting in 35,328 transformants. Mutant cells were plated at limiting dilution in 96 well dishes to favour clonal growth. After growth for 1-2 days, wells containing a single mutant were transferred into 368 master plates. This was replicated to generate plates for DNA extraction (to identify insertion points) and to provide stocks for mutant strain storage.

**Figure 3.**
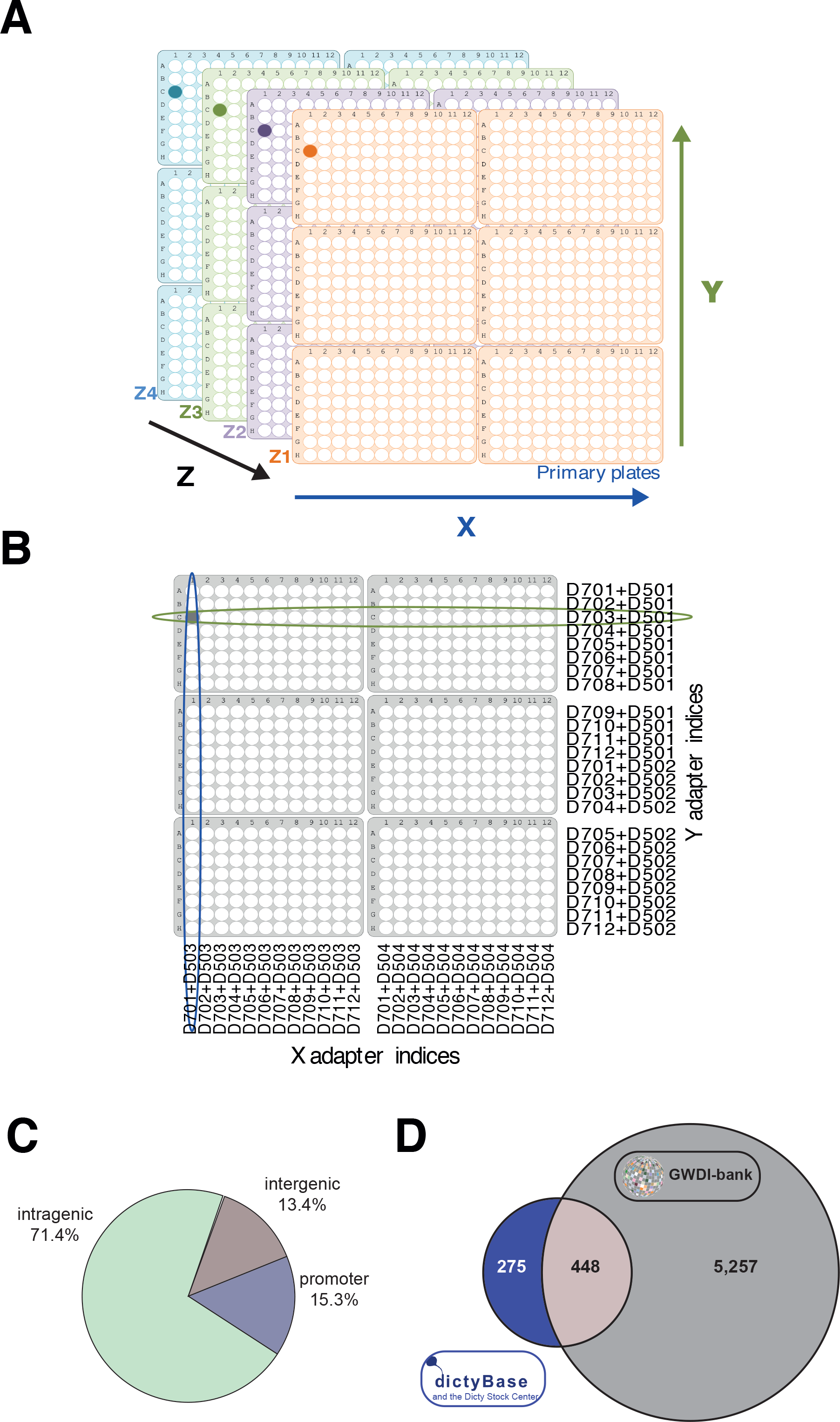
Gridding and multiplexing of clonal mutants for sequencing. **A. Master grid contains 2304 wells.** Mutants made with one of four GWDI barcodes were grown up clonally in 96-well plates. A grid containing six plates of mutants with the same barcode was generated. This was repeated for mutants with each GWDI barcode to make a stack of four grids. **B. Multiplexing of mutants for sequencing.** Cells in each row (X), column (Y) and stack (Z) were pooled and gDNA prepared, to make a grid that is compressed into the X and Y coordinates. Unique combinations of X and Y barcoded sequencing adapters were ligated to each gDNA preparation before Illumina sequencing. The intersection of common mutants provides the well location for each mutant. The four mutants present in the compressed grid can be separated to stacking grids by their unique Z-barcode. **C. The collection of individual *Dictyostelium* REMI mutants** The REMI-seq mutant resource consists of 21,529 individual mutants. The majority of these mutations are in coding sequence or within putative promoter sequences and thus likely to result in gene disruptions. The REMI-seq resource consists of mutations in 5705 genes, resulting in >7.5 fold increase in the number of mutants available. The catalogue of mutants can be found on the REMI-seq website (remi-seq.org) and individual mutants can be ordered from the Dicty stock centre.

Mutant identification was based on a conceptual 24×24 grid, the equivalent of a 2 × 3 array of 96 well plates (Figure 3A). Mutants from each row and each column were pooled and their DNA extracted (Figure 3A). In this way, each mutant is represented twice, once in a row (X) and once in a column (Y), and can be identified by coordinating its XY position within the 24 × 24 notional grid. To do this, a series of 16 library adapters (D501-D504 and D701-D712) required for Illumina sequencing were made that introduce a unique 8bp barcode to each fragment terminus (Figure 3B). A further grid dimension was added with a Z coordinate, corresponding to a notional stack. Each layer of the stack was identified by using a different insertion fragment, again each marked by a unique barcode (Supplementary File 4). After DNA sequencing, the 3D-grid position of each mutant can be inferred computationally by its unique XYZ coordinate (Figure 3A & B). Consequently, each sequencing run had the capacity to generate information from 4,608 wells or mutants (see https://github.com/NicoleGruenheit/grid_analyser for information about the layout and analysis).

DNA sequences, both upstream and downstream, of all integration sites were identified by first filtering the read files for the presence of short sequences derived from the insertion fragment and then aligned to the *Dictyostelium* AX4 reference genome (Supplementary File 3). We were able to assign 21,529 independent mutants, corresponding to 12,247 different genomic insertion sites (Supplementary File 5 and www.remi-seq.org). We termed this resource the GWDI (pronounced “goody”) bank. Our *in silico* genome analyses predict that most genes should be accessible to REMI-seq. Using a window size of 40,000 bp, which divides the 35 Mbp genome into 861 non-overlapping windows, we compared the number of possible sites per window to the number of observed insertion points in the pool (see methods). Out of these 861 windows, only 15 contained less (coldspots) and 28 contained more insertion points (hotspots) than you would expect under a uniform distribution using an adjusted *p*-value cut-off of 0.05. Therefore, even though sites in essential genes are likely refractory to REMI mutagenesis, the overall frequency with which different sites are hit is close to random. Indeed, regions associated with hotspots only account for 8% of the observed insertions in the pool. Consequently, we were able to catalogue 12,247 different insertion points, of which 8,739 (71%) were found to lie within coding regions of genes, whilst another 1,872 were found within putative promoter regions of genes (defined as <500 bp upstream of the start codon) and thus also likely to disrupt gene function (Figure 3C). In total, our REMI-seq approach has resulted in the production of a collection of mutants in 5,704 different genes (ie 43% of the genome). They are catalogued and searchable (www.remi-seq.org). They have also been deposited in the *Dictyostelium* stock centre and are freely available on request (Figure 3C).

### Use of GWDI-bank in functional genomics pipelines

Direct access to a bank of mutants encompassing nearly 6,000 *Dictyostelium* protein coding genes offers a major interface with other functional genomic resources and high-content data collections. A number of other standard ‘omics’ approaches are available for *Dictyostelium* research, such as transcriptomics and proteomics. GWDI bank, generated by REMI-seq, has the capacity to integrate across these modalities by connecting genotype to phenotype. To explore this potential, we undertook a pilot experiment based on the proteomics study of actin cytoskeletal associated proteins of Sobczyk et al (2014) (Sobczyk, Wang and Weijer, 2014). This study used SILAC to identify over 470 proteins that associate with the actin cytoskeleton during chemotaxis. The challenge is to identify those proteins that are regulatory and important for actin cytoskeletal dynamics, and this requires the generation of a mutant in each gene. We, therefore, selected 12 of the proteins identified in this analysis represented by 2 or more independent mutants in the GWDI bank, of which the function of five of proteins was unknown (Supplementary File 6).

*Dictyostelium* cell motility is driven by F-actin polymerisation, and changes in its regulation are likely to translate into altered cell movement. Almost all *Dictyostelium* cell motility assays are based on starved mid-aggregation stage cells undergoing chemotaxis to cAMP or chemotaxis of bacterially grown-cells to folate. Developed cells show a significantly higher speed and directionality of movement compared to those in growth phase (Varnum, Edwards and Soll, 1986; Alonso, Stange and Beta, 2018). This difference may arise due to the influence of different molecular pathways controlling chemotaxis (Nichols, Veltman and Kay, 2015). However, screening for chemotaxis defects in large numbers of mutants is difficult. We therefore devised a simple assay for the motility of growing cells under normal growth conditions. In our assay, cells were plated into a 96-well microtitre plate at a low enough density to allow cell movement to be recorded by video-microscopy. Once plated and adhered to the plate surface, individual cell positions were recorded every 2 minutes for a minimum of 44 minutes using high content microscopy. The distance travelled between time points was measured, and when summed gives the total distance travelled. This assay allows each mutant to be assayed under identical assay conditions and its ease of set up offers a future basis for high throughput screening (HTS).

The motility of the 12 mutants identified through proteomic analysis and selected from the GWDI bank was assayed. Although individual cells in culture show substantial variation in distance travelled, significant differences were found for some mutants when compared to wild the cells. For example, the DOCK protein-family gene *docA* has been examined in detail previously (Sobczyk et al., 2014), and found to affect chemotaxis in starved cells. Similarly, we found that it affects motility in slow moving growth phase cells. We also found ziziminA (zizA), another DOCK family protein shown to localise to the microtubule organising centre (MTOC), but with no reported phenotype (Pakes *et al.*, 2012) has elevated motility in this assay. Finally, we tested another DOCK protein *docB*, which is represented as a single mutant cell strain in GWDI bank (not shown), and also found it to exhibit increased motility. In contrast, we found that *elmoE* showed reduced motility, as did *roco10* and *rapgap1*. The ELMO family of proteins forms a complex with DOCK proteins to function as Rac-GEF. In *Dictyostelium* chemotaxis, loss of *elmoE* reduces cell speed and shows reduced activation in response to cAMP stimulation (Yan *et al.*, 2012). Loss of *rapgap1* is also known to affect cell motility deficits during chemotaxis, and here we show that this may be the case for non-chemotactic growth phase cells (Jeon *et al.*, 2007). In addition, our assay showed that previously untested gene mutations of roco10, DDB_G0289829 and DDB_G0277675 all reduced cell motility, while DDB_G0277997 encoding a protein of unknown function elevated motility. This pilot experiment thus demonstrates that a simple phenotype can be employed to screen GWDI-bank, and connect the output of proteomic studies to gene function.

### REMI-seq can monitor mutant population dynamics for parallel phenotyping

‘Parallel phenotype’ approaches provide a powerful method to determine the relationship between genotype and phenotype (Wang *et al.*, 2014; van Opijnen, Lazinski and Camilli, 2015), by establishing mutants that confer a selective advantage or disadvantage within the population under different conditions. We thus assessed the ability of REMI-seq to monitor the relative abundance of each mutant in the population. We first sequenced a test population, estimated to contain approximately 20,000 mutants, and found we could detect 17,812 independent REMI insertions (Supplementary File 7). Analysis of vector specific barcodes revealed 8,593 mutant insertions to be present only once, with 4,020 and 393 insertions arising from two and three independent transformations respectively. This pooled population contains mutations in the coding sequence of 5,268 different genes (44% of all genes in the genome), or 5,811 genes when promoter insertions were included. Of these, independent alleles were identified in 3,501 genes, with 898 of these genes represented by different insertion points within the same gene, 1,077 genes with at least one identical independent insertion at the same locus, and 1,526 with both.

To test technical reproducibility, the population was split into triplicate samples and sequenced. Read counts for each insertion site were highly correlated (*r* = 0.92; *r* = 0.92; *r* = 0.91, *p* < 2.2 × 10^−16^ for all comparisons) (Figure 5A), demonstrating a good overall quantitative reproducibility. We next probed sources of variability, and established that this is mainly determined by mutant frequency in the population, and the linked tag sequence read-depth. This was particularly prominent for mutants with read counts <100. When each pair of technical replicates was compared, up to 78% of these low abundance mutants with <100 read counts in the starting library was missing from, or ‘dropped out’ of, at least one replicate (Figure 4B). However, the drop-out rate decreased dramatically when read counts were > 100, and was zero when read counts were >1000 (Figure 5B).

**Figure 4.**
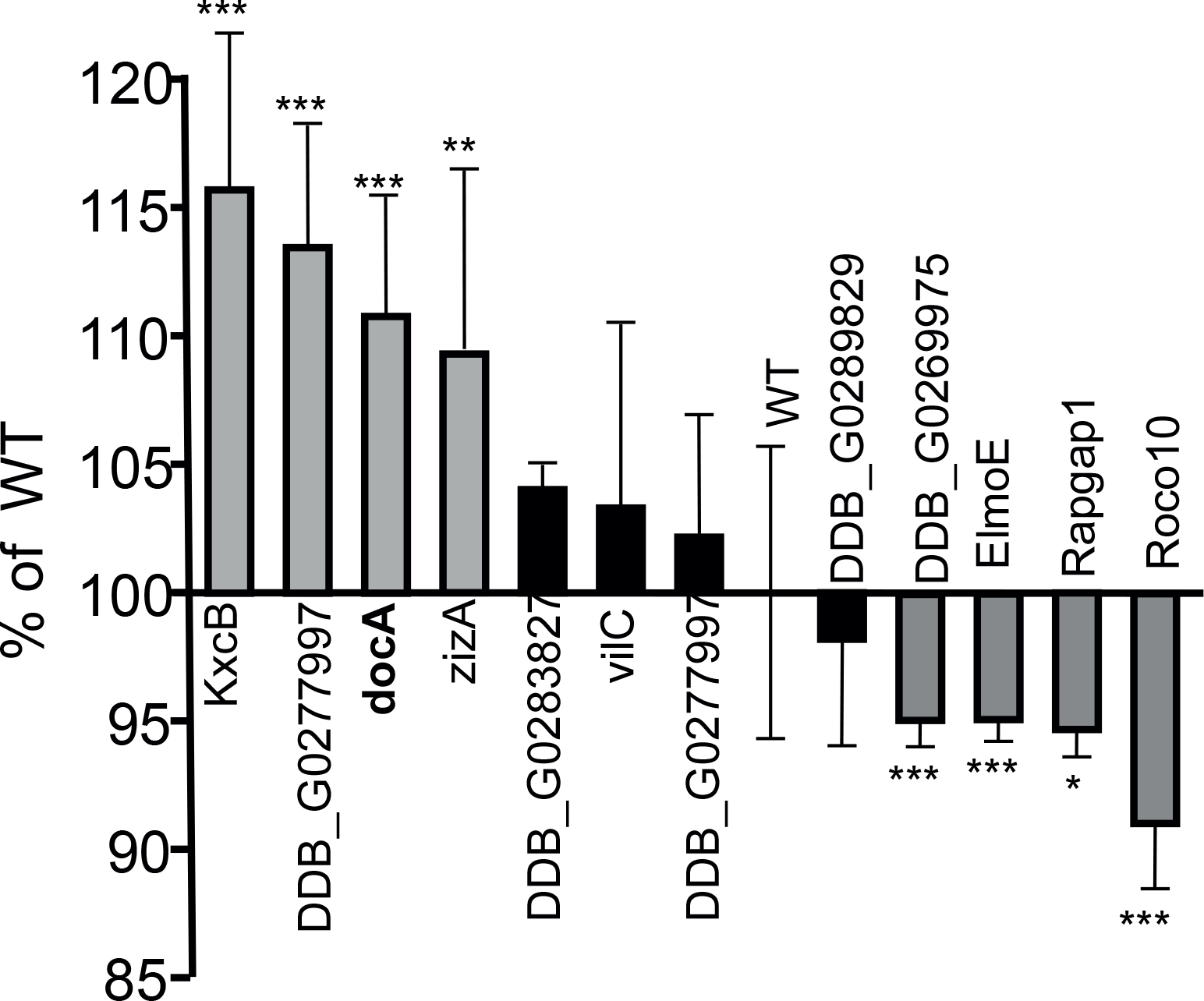
Screening mutant strains for motility changes in growth phase culture. Cells from two independent strains for each mutant were plated in wells of a 96-well plate in parallel cultures. 100-1000 cells were imaged for 44 minutes. Histogram shows the mean the Total Path Length for the two strains of each mutant, expressed as a percentage of the wild type control (WT), error bars show standard deviation. * = p<0.05, ** = p<0.005 and *** p<0.001

**Figure 5.**
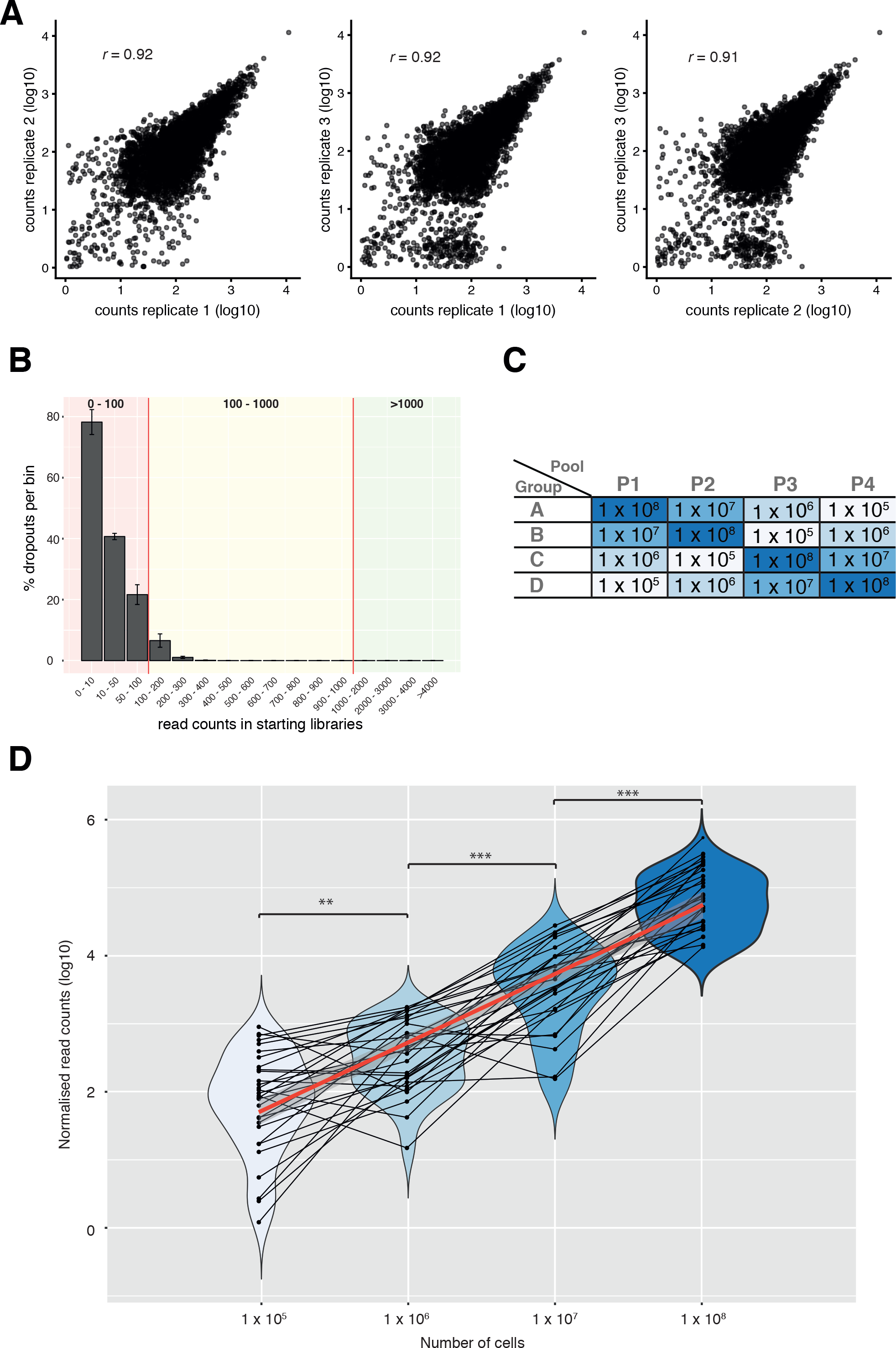
REMI-seq can be used to quantify the relative abundance of mutants. **A. REMI-seq exhibits high technical reproducibility.** Three preparations of gDNA from a pool of ^~^20,000 mutants were sequenced. Normalised read counts for each mutant are highly correlated across each replicate. **B. Technical reproducibility and dropout rate is dependent on read count.** Mutants were divided into bins dependent on the average starting read count between each two replicates of the starting library. The percentage of mutants detectable in only one library was calculated for each bin. Error bars represent the average of three pairwise comparisons. This revealed that technical dropouts are highest in mutants with very few cells in the starting library, which then results in very few extracted tags and low read counts (< 100). **C. Schematic of experiment to determine quantitative dynamic range of REMI-seq.** Cells of 32 defined mutants were distributed into four groups (A, B, C, D) of eight mutants each (see Supplementary file 8). 1 × 10^5^, 1 × 10^6^, 1 × 10^7^ or 1 × 10^8^ mutant cells from each group were mixed to form four different Pools (1-4). **D. REMI-seq can be used to quantify mutant abundance over >100 fold range.** Pools of mutants spiked at different frequencies were subjected to the REMI-seq protocol and sequenced. Tags were extracted from the sequencing files, normalised to the number of reads per pool and identified positions compared to the list of defined mutants. A violin plot for each density was derived from the logged normalised read counts for each mutant. The number of cells was highly correlated to the normalised read counts (*r* = 0.55, p = 2.439 × 10^−11^). All means are significantly different to each other (one-sided t-test, 1 × 10^5^ vs 1 × 10^6^: 0.002; 1 × 10^6^ vs 1 × 10^7^: 0.0002; 1 × 10^5^ vs 1 × 10^6^: 0.002; 1 × 10^7^ vs 1 × 10^8^: 0.0002). See Supplementary file 8 for separate results of each group of mutants.

We next tested the sensitivity and quantitative range of our population REMI-seq method. Four groups of mutants were generated (A, B, C, and D), with each group composed of eight mutants with known insertion points (Supplementary File 8). Each group was mixed together at different frequencies across three orders of magnitude to generate pools 1-4 (Figure 5C). This experimental design allowed us to determine the variation in detection sensitivity between different insertion points spiked at identical frequencies. Furthermore, it allowed us to determine the quantitative dynamic range of each mutant when mixed at different frequencies across the four samples. Overall, these data revealed that input and output frequencies of all mutants were highly correlated (*r* = 0.55, p = 2.4 × 10^−11^) and tracked in frequency across the different groups as expected (Figure 5D). It should be noted, however, that the standard deviation of the read counts, i.e. the noise in the data, derived from the lowest input frequency is higher than other samples. This is because these mutants had read counts of approximately 100, highlighting the importance of treating these mutants separately. Together, these data illustrate that by dividing the mutants into different read count bins, carrying out all experiments with at least two biological replicates, and ensuring that there is sufficient sequencing depth, it is possible to use REMI-seq to identify mutants that decrease (or increase) in frequency within the population, reflecting altered selective pressures.

### Population dynamics of REMI-seq mutants under differential growth conditions

To pilot REMI-seq for tracking population dynamics and mutant fitness, our pooled population of 17,812 mutants was selected for growth either on the standard bacterial laboratory food source *K. aerogenes* or under axenic conditions in liquid HL-5 media. Growth on bacteria requires phagocytosis of large particles (King *et al.*, 2013) whereas growth in liquid medium requires micropinocytosis of fluid (Veltman *et al.*, 2016; Bloomfield *et al.*, 2015), and therefore a different spectrum of genes are likely to underpin each mechanism. The population was serially passaged in duplicate through growth either in HL-5 axenic culture for up to 72 generations or in association with *K. aerogenes* over 200 generations (Figure 6A). The samples were sequenced at 24, 48, and 72 generations of axenic growth or 100 and 200 generations of growth on *K. aerogenes*. As expected, the read counts of each biological replicate were highly correlated (*p* = <10^−15^, Supplementary figure 1), and the amount of technical variation was dependent on the initial read counts (Supplementary figure 1). To determine if mutants became under- or overrepresented, we compared the relative read count of each mutant in the starting library to the relative read count after growth selection. Mutants were first separated into the three bins (1-100, 100-1000, >1000 reads) according to their read count in the starting library. This allowed us to determine the average change in behaviour of each mutant within each bin to identify those mutants whose abundance deviated significantly (Z-score) from this behaviour (Figure 6B and Supplementary figure 2). This takes into account that mutants with increased fitness will increase in frequency and alter the relative abundance of neutral mutants (see Supplementary figure 1). This is especially evident in the stronger axenic growth selection, where the number of reads for neutral mutants decreases substantially with many disappearing altogether (Supplementary figure 1). In addition, mutants with initial read counts of <100 that decreased in frequency were discarded from the analysis because the technical dropout rate in this group is very high (Figure 5B and Supplementary file 6). Together, these steps allowed the identification of mutations that significantly affect fitness during the selection.

**Figure 6.**
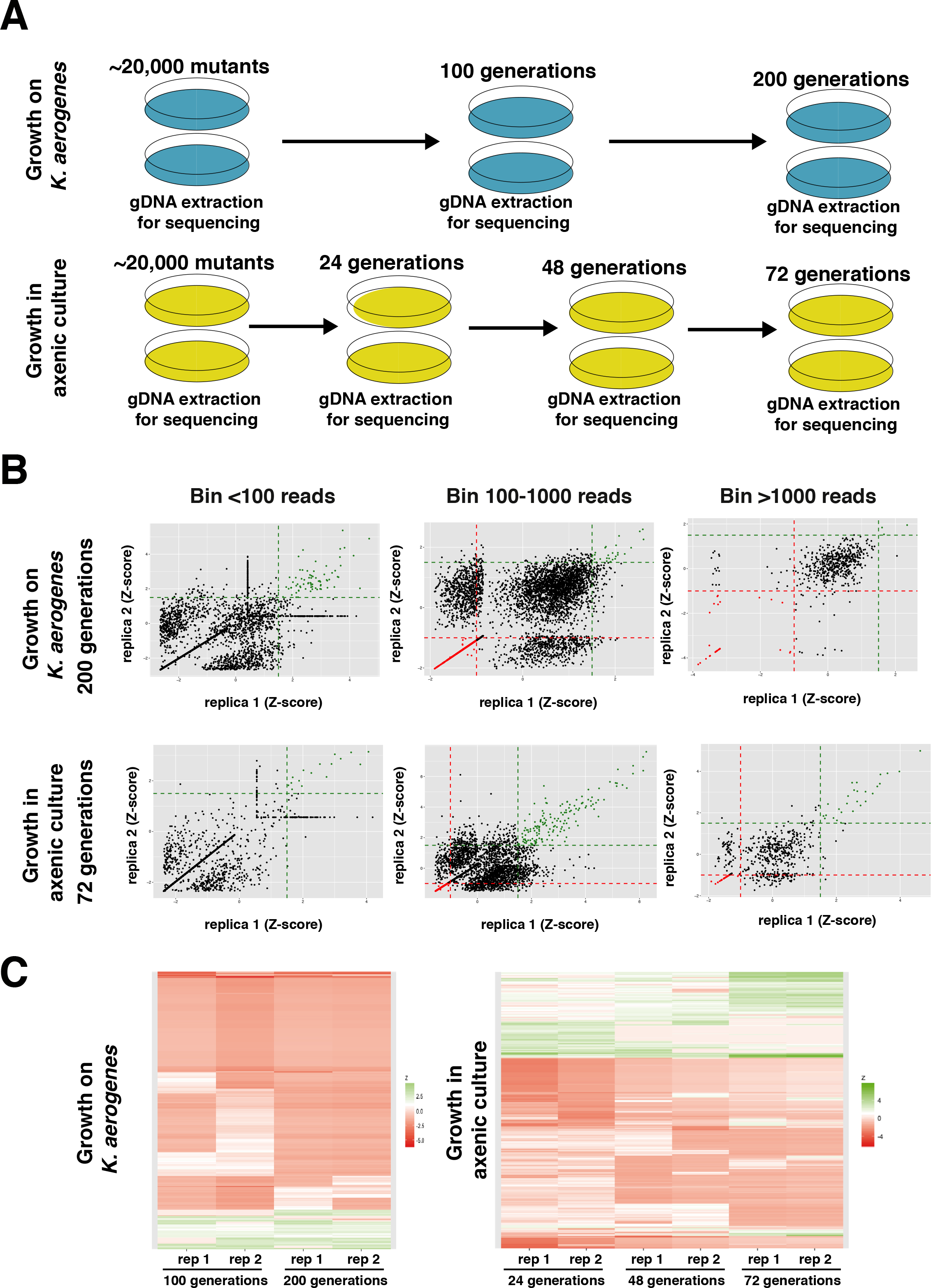
REMI-seq parallel phenotyping to identify mutants that affect axenic or bacterial growth. **A. Schematic of selections to identify mutants with growth advantages and disadvantages.**A pool of ^~^20,000 mutants was plated in duplicate in HL5 medium for axenic growth, or in association with *Klebsiella aerogenes* bacteria. Cells were grown for 24 (axenic) or 100 generations (Ka) before a sample was harvested for sequencing. The remaining cells were diluted and grown up for a further 24 (axenic) or 100 (Ka) generations before a second sample was taken for sequencing. Axenic cells were diluted and grown up for a further 24 generations before sequencing. **B. Identification of mutants with growth advantages and disadvantages**. The abundance (read count) at each round of each mutant was compared to the start pool. Mutants were first divided into bins based on their read count in the start pool in order to identify mutants that deviated in abundance (Z-score) significantly from other mutants with similar read counts. Mutants with a Z-score of >1.5 (green dashed line) in both replicates were considered to have increased in abundance. Mutants with a Z-score of <−1.0 (red dashed line) were considered to have decreased. In bin <100, the variation due to technical dropouts resulted in a high false discovery rate, and mutants that decreased were not considered. Replicas are highly correlated. Results are shown at the end point of the selection (after 200 generations of Ka growth and 72 generations of axenic growth). Data from other rounds showed similar patterns (see Supplementary figure 2). **C. Mutant behaviour is consistent across rounds of selection.** Mutants were identified with a significant advantage or disadvantage at each stage of the selection. The level of enrichment or depletion (deviation from expected or Z-score) was plotted for each mutant at all rounds.

Our analyses revealed that a significant proportion of mutants that are enriched or depleted in one round of the selection, were often also enriched or depleted in future rounds (Figure 6C). As expected, few mutants switched from being significantly enriched to depleted, or vice versa. Some mutants, however, were only enriched or depleted in early or late rounds of the selection. This is likely due the complex dynamics of selection. For example, weakly advantageous mutations may be lost in later rounds as stronger mutants take over the population. Alternatively, the same mutants could take several rounds to become significantly enriched. Similarly, it is sometimes impossible to determine if mutants are significantly depleted in later rounds if all mutants in the bin as a whole have essentially been removed (i.e. things cannot decrease below zero). These data not only highlight the quantitative reproducibility of the REMI-seq technique, but also show the importance of observing multiple rounds.

To confirm that the gene mutations identified from parallel phenotyping do indeed confer positive or negative fitness under the different growth conditions, we tested independently generated mutations at the same locus isolated from the gridded GWDI-bank. The fitness of each mutant was validated in 1:1 competition with their parental strain (Figure 7A). Consistent with expectations, fitness in 1:1 competition was highly correlated with fitness measured in the equivalent population screen (Figure 7B and Supplementary figure 3). Together these data demonstrate that REMI-seq provides an effective method to assign phenotype to genotype in *D. discoideum*, as well as suggesting that the fitness effects observed here did not arise as a consequence of interference competition. Finally, it shows how integration of the gridded mutant set with pools used for parallel phenotype analysis provides an effective pipeline for functional genomic analyses in *D. discoideum*.

**Figure 7.**
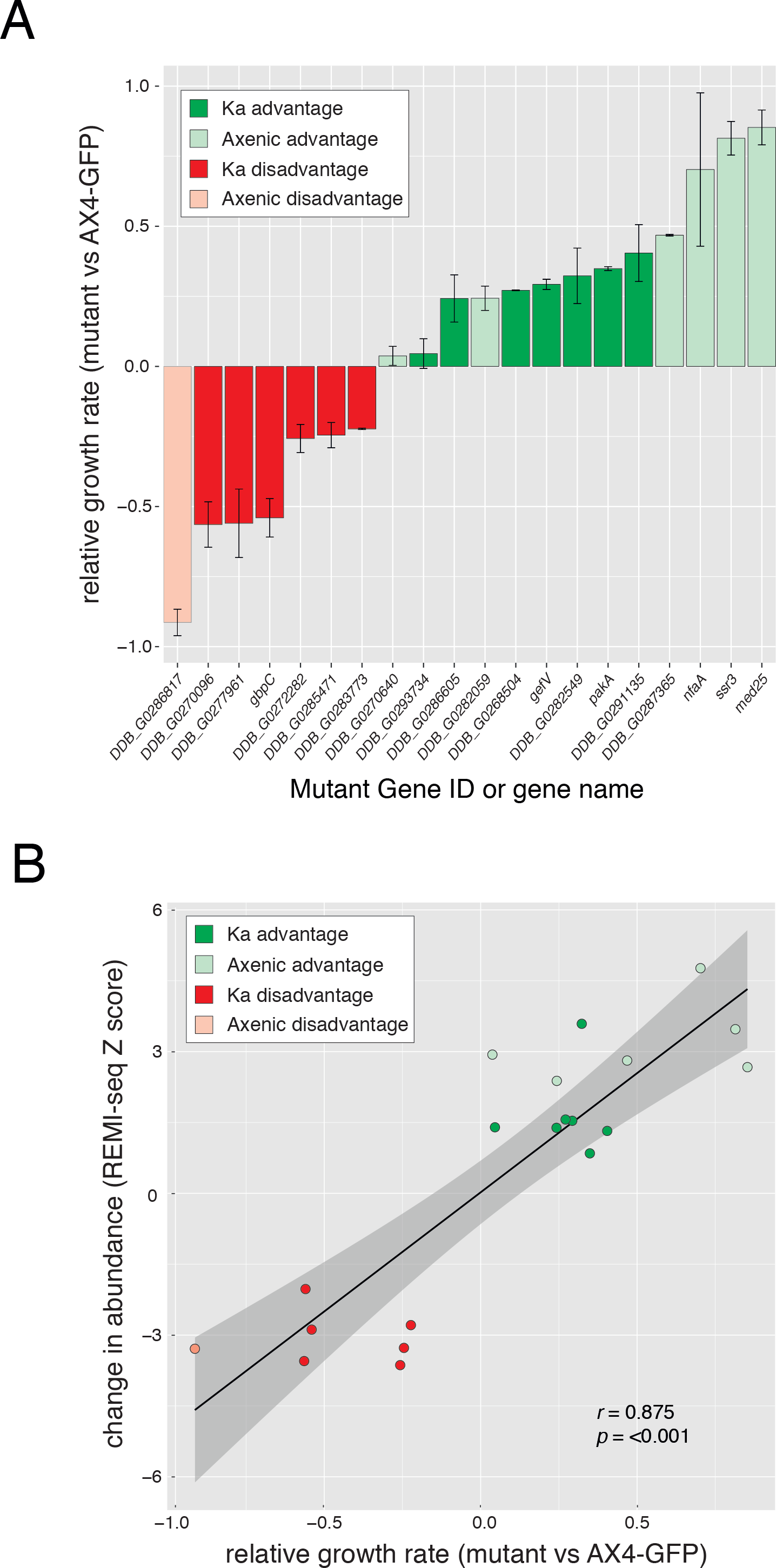
Validation of mutants identified by REMI-seq. **A. Mutants identified by REMI-seq exhibit growth advantages or disadvantages.** Independent isolates of 20 mutants identified by REMI-seq were tested for growth effects in competition with the parental AX4 strain. 18 mutants showed significant differences, and all mutants identified by REMI-seq as disadvantaged grew more slowly than wild type. **B. REMI-seq can quantitatively predict fitness effects in growth competition.** There is a strong correlation between the experimentally measured growth rate of each mutant and the relative change in abundance of each mutant inferred by REMI-seq at the final round of selection. Similar results can be seen when the data from every round is compared to the growth rate (see Supplementary figure 3).

### Functional genomics of axenic and bacterial growth

Many functional genomic approaches infer gene function from changes in transcript abundance. However, the relationship between gene function and gene expression may be complex, and non-linear. For example, studies in *S. cerevisiae* (Winzeler *et al.*, 1999) have revealed a surprising lack of correlation between differential transcriptional regulation and a requirement for fitness under those conditions, where fewer than 7% of upregulated genes were also required for optimal growth (Giaever *et al.*, 2002). Parallel phenotyping using REMI-seq provides a method to directly address gene function in the absence of other information. To investigate how this may relate to gene expression, we compared the transcriptome of cells after two days of growth on bacteria or in axenic medium, with the gene set identified by REMI-seq. However, we found no correlation between differential gene expression and growth fitness of mutants with insertions within the coding sequences or promoter regions (up to 500 bp upstream of the transcription start site) of genes (Figure 8A).

**Figure 8.**
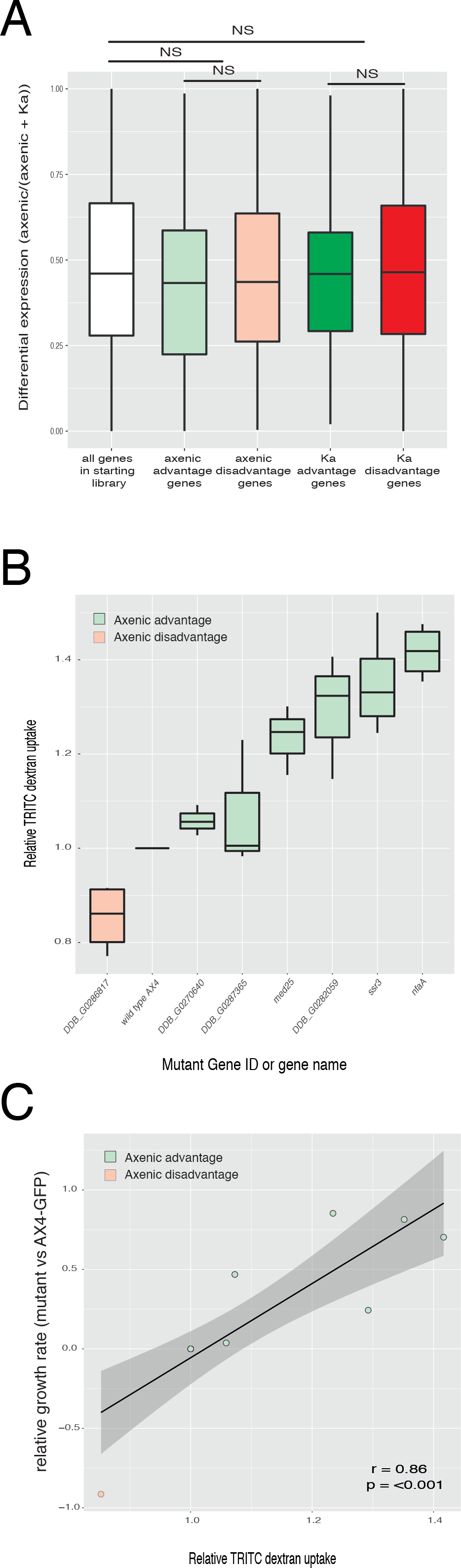
REMI-seq identifies novel regulators of axenic and bacterial growth. **A. Differential expression does not correlate with gene requirement.** Gene expression was compared in cells grown in axenic medium or in association with Klebsiella aerogenes. Differential expression was calculated as an expression index where a value of 1 represents expression exclusively in axenic growth and a value of 0 represent expression solely in bacterial growth. The expression index of genes in which mutations result in increased or decreased growth rate in axenic or bacterial growth was compared to all other mutations in the library that could be ascribed to the likely functional disruption of a single gene. No significant differences were found. **B. The rate of macropinocytosis is affected in mutants with altered growth in axenic medium.** The rate of TRITC-dextran uptake was compared in a selection of seven mutants identified in the axenic growth selection to parental AX4 wild type cells. The rate of uptake was significantly different to wild type in six mutants. **C. The rate of macropinocytosis correlates with growth rate in axenic medium.** Growth rate in completion with wild type cells (see Figure 6A) was compared to the rate of TRITC-dextran uptake.

One reason for this lack of correlation could be that differentially expressed genes (DEGs) are actually neutral when mutated. Alternatively, if mutations in these genes result in a strong growth defect under either condition, they could be lost from the REMI-seq starting library before the start of the selection. This is because some growth in HL5 or Ka is actually required for the generation and sequencing of the REMI-seq starting library. Consistent with this latter idea, GO terms associated with the identified DEGs were actually underrepresented in the REMI-seq starting library (Supplementary file 7), including genes required for the core machineries of import into the cell, vesicle mediated transport and ribonucleoprotein complex assembly. In addition, any remaining mutants associated with these GO terms tended to have lower than average read counts in the starting library (Supplementary file 6). These data suggest that for a subset of genes, there is good agreement between gene expression and gene function.

We next further analysed genes that were identified because they displayed a phenotype in the REMI-seq screen, but showed little correlation to the degree of differential expression. These genes are interesting because this would have precluded their identification in transcriptomic studies. Genes that affected growth on bacteria were strongly enriched for processes associated with cellular responses to stress, especially the response to oxidative stress (Supplementary file 7). These included redox homeostasis and superoxide removal (including thioredoxins, glutaredoxin, and protein disulphide isomerase encoding genes), DNA repair, pteridine production and copper ion transport. The role of ROS production in bacterial growth and killing has been unclear (Dunn *et al.*, 2017). However, these data suggest the ability of *Dictyostelium* cells to withstand oxidative stress is crucial for normal growth. It is also interesting to note the enrichment of terms associated with lipid composition, including genes associated with sphingolipid metabolism. Indeed, it has recently been suggested that lipid composition may play a key role in host pathogen interactions in macrophages (Teng, Ang and Guan, 2017), but so far has not been noted in *Dictyostelium*. These studies thus highlight the importance of processes such as prey recognition, killing and toxin tolerance in addition to uptake for bacterial growth.

Axenic growth requires cells to take up large volumes of fluid from the surrounding environment by micropinocytosis. Indeed GO terms associated with fluid uptake are enriched in genes identified through differential gene expression analysis, and are underrepresented in the REMI-seq starting library. However, we also found that selection for growth in axenic medium resulted in the identification of mutants that were enriched for GO terms associated with signalling, especially proteins that affect phosphorylation such as protein kinases and histidine kinase response regulators (Supplementary file 9). Furthermore, we found enrichment of mutations in genes encoding putative regulators of small GTPase activity, including *nfaA* (which also encodes a Ras-GAP) and DDB_G0286817 (which encodes a putative regulator of GAP activity). These findings are consistent with the idea that the rate of micropinocytosis can be regulated by environmental cues, such as nutrients or amino acid availability (Williams and Kay, 2018b) or by changes in Ras activity (Bloomfield *et al.*, 2015). We, therefore, tested whether mutants that show increased or decreased rates in growth in HL5 medium, also affect the rate of macropinocytosis. The rate of TRITC dextran uptake was significantly different to wild type in six out of seven mutants tested (Figure 8B). Furthermore, this correlated strongly with the growth rate of each mutant in HL5 medium (Figure 8C). Together, these studies highlight the effectiveness of REMI-seq for identifying key regulators of biological processes. Furthermore, these data highlight the importance of taking parallel approaches to ascribe function to certain genes or highlight processes, which may otherwise be missed if gene expression analyses are considered in isolation. Because their identification is often refractive to identification by other methods, REMI-seq provides a step change towards the goal of defining the genetic ‘components list’ underlying cell and developmental systems.

## Materials and Methods

### Generating the insert DNA

The pGWDI plasmids were derived from pLPBLP (Faix *et al.*, 2004). Double stranded oligonuclotides were incorporated at two positions within pLPBLP. AJBgwdi001-F and AJBgwdi002-R were incorporated between the *Kpn*I and *Hind*III sites; AJBgwdi003-F and AJBgwdi004-R were incorporated between the *Pst*I and *Spe*I sites. iLB and iRB are vector specific indexes (see Supplementary Table 1). Lowercase text denotes sticky ends for ligation. Sanger sequencing confirmed cloning was successful.

AJBgwdi001-F: 5’-CGGATCCGTTGGACTGCTG[iLB]TAGGGATAACAGGGTAATA-3’
AJBgwdi002-R: 5’-agctTATTACCCTGTTATCCCTA[rc-iLB]CAGCAGTCCAACGGATCCGgtac-3’
AJBgwdi003-F: 5’-GATTACCCTGTTATCCCTA[iRB]CAGCAGTCCAACGGATCCA-3’
AJBgwdi004-R: 5’-CTAGTGGATCCGTTGGACTGCTG[rc-iRB]TAGGGATAACAGGGTAATCTGCA-3’

There are 8 versions of pGWDI: pGWDI-G1, -G2, -G3, -C4, -G5, -C6, -C7 and -C8. The resulting GWDI-<u>G</u> inserts, that are derived from the plasmids by amplifying a 1,593 bp region around the blasticidin resistance cassette using the T3 and T7 primers, have GATC-sticky ends generated by BamHI cleavage, whereas the GWDI-<u>C</u> inserts have -CATG sticky ends generated by SphI cleavage. Each arm of each insert can be distinguished by unique 6 bp indices (Supplementary Table 2) and contains Isce-I and MmeI recognition sequences that are necessary to identify the location of the insert within the genome. The arms terminate in 4 base overhangs.

### REMI mutagenesis

*Dictyostelium* AX4 cells, strain I.D. DBS0235554, were grown in shaking suspension in HL-5 medium including glucose (Formedium) with PVS at 22°C until they reached mid-log (0.5-1.0 ×10^7^ cells/ml). Unless otherwise stated all steps are conducted on wet ice. Cells (1 ×10^8^) were cooled for 15 minutes before collection by centrifugation for 2 minutes at 540g at 4°C. Cells were washed twice with cold KK2 + sucrose (16.1mM KH2PO4, 3.7mM K2HPO4, 50mM sucrose) and collected in the same manner. Cells were re-suspended in KK2+sucrose to a density of 1.25×10^7^ cells/ml. Insert DNA (70 μg) and 7 ml cells were combined and incubated on ice. After 5 min, 6.6 U enzyme was added and gently mixed. Aliquots (800 μl) were transferred to 8 pre-cooled 4 mm gap-width electroporation cuvettes. Cells were transformed using an ECM399 electroporation generator set to 900 V. The ECM399 has a 150 Ω resistor and a 36 μF capacitor. The output voltage was typically 810 V and the time constant was ^~^2 ms. CaMg (8 μl of 100 mM CaCl_2_ and 100 mM MgCl_2_) was immediately added to the cells following electroporation and they were transferred gently to 500 ml of rich media (filter sterilised HL-5 medium including glucose, 10% horse serum, 1x PVS. 100x PVS comprises 3g penicillin G, 5g streptomycin sulphate, 10 mg folic acid, 30 mg vitamin B12 in 500 ml H_2_O, filter sterilized and stored at 4°C in the dark. This was repeated for the seven remaining aliquots. Cells were transferred to 96 wells (100 μl per well) using an Integra VIAFLO 96 electronic pipette and incubated at 22°C overnight. Transformants were selected by the addition of blasticidin to a final concentration of 10 μg/ml the following day.

### Preparation of target for Illumina Sequencing

Genomic DNA was collected from cells grown in association with *K. aerogenes* on SM agar plates. Nuclei were collected from approximately 5 × 10^8^ cells that had been washed at least 5 times in cold KK2 (16.1mM KH2PO4, 3.7mM K2HPO4) and then resuspended in 30 ml nuclei buffer (40 mM Tris, pH 7.8, 1.5% sucrose, 0.1 mM EDTA, 6 mM MgCl2, 40 mM KCl, 0.4% NP-40 substitute, 5 mM DTT) followed by centrifugation at 4,000 g for 30 min, 4°C. Supernatant was discarded.

Pellets were suspended in EDTA to a final volume of 100 μl and a final concentration of 100 mM, before 100 μl of 10% sodium lauryl sarcosyl was added, mixed and incubated at 55°C for 20 min. 4M ammonium acetate (250 μl) was added prior to centrifugation at 20,000 g for 15 min, 4°C. One volume of supernatant was added to 2 volumes of absolute ethanol and mixed well. The samples were centrifuged at 20,000 g for 10 min, 4°C, and supernatants were then discarded. Pellets were washed with 300 μl of 70% ethanol, dried and suspended thoroughly in 50 μl of 10 mM Tris pH 7.8, containing RNase A and RNase T1 (10 U/ml and 400 U/ml respectively, Ambion). Samples were centrifuged at maximum force for 3 min to collect any insoluble material and supernatant was transferred to a clean microcentrifuge tube. The integrity of the samples was visualised by separation by electrophoresis through a 1% agarose gel.

MmeI and I-SceI were used to excise the DNA of interest, i.e. the genomic DNA – insert DNA junction. The insert contains an I-SceI recognition sequence; this sequence does not appear elsewhere in the genome. The insert also contains an MmeI recognition sequence; MmeI cuts the DNA 19/20 bp from its recognition sequence, and was positioned so that digestion captured 19/20 bp of gDNA.

MmeI (10 U) was used to digest genomic DNA (0.5 – 1 μg) overnight in a total reaction volume of 200 μl (NEB). MmeI was inactivated by incubation at 65°C for 20 min. The samples were further digested for 6 hr by 25 U of I-SceI (NEB). I-SceI was also heat inactivated. DNA was collected by ethanol precipitation using standard procedure and suspended in 1x T4 DNA ligation buffer.

Indexed adapters (D7 and D5) were ligated to the digested DNA. Different combinations of D7 and D5 indices were used to tag each sample. D7 adapters have a 4-bp ‘-TTAT’ 3’ overhang that is complementary to the overhang generated by I-SceI cleavage. Likewise, D5 adapters have a 2-bp ‘-NN’ 3’ overhang, which is complementary to overhangs cleaved by MmeI. Ligation reactions (200 μl) comprised digested DNA, 1x T4 ligation buffer (NEB), D7 and D5 pre-annealed indexed adapters (2 ng and 200 ng, respectively) and T4 DNA ligase (400 U, NEB), and were incubated at 16°C overnight. See below for adapter preparation.

In some cases the abundant mitochondrial large subunit ribosomal RNA (rnlA) impeded processing. Thus, samples were digested with 10 U *psh*AI in 1x CutSmart buffer at 37°C for 3 or more hours (NEB).

Amplification of the DNA of interest was achieved by PCR using primers specific to the D7 and D5 adapters. PCR was conducted using GoTaq® G2 Flexi DNA Polymerase (1.25 U, Promega), the P5 primer (0.5 μM, 5’-AATGATACGGCGACCACCGA-3’), the applicable D7_a single oligonucleotide (0.5 μM), 1x flexi buffer, 1 mM MgCl_2_, 0.8 mM dNTPs and sample (^~^ 5 ng genomic DNA). The target was amplified using the following conditions: 2 min initial denaturation at 95°C, 35 cycles of 95°C for 30 sec, 66 for 5 sec, 68°C for 1 min; and a final extension for 10 min at 68°C.

Samples were separated by agarose gel electrophoresis (3% gel) to confirm the presence of a band of the expected size (183 bp). Each sample was processed individually until this point. According to the band intensity observed via agarose gel electrophoresis samples were combined in equal molar ratios.

Size selection was used to enrich the DNA of interest within the pooled sample. The pooled sample was separated by electrophoresis using a 3.5% low melting temperature agarose gel (NuSieve™ GTG™ Agarose, Lonza). The band corresponding to approximately 183 bp was excised and DNA was extracted (QIAquick Gel Extraction Kit, Qiagen). The resulting DNA was size selected a second time in the same manner with the exception that DNA was extracted from the agarose gel using the Qiagen MinElute® Gel Extraction Kit.

The resulting sample was analysed using a D1000 ScreenTape and Agilent 2200 TapeStation system (Agilent Technologies), a Qubit® 3.0 Fluorimeter, and sequenced using an Illumina MiSeq with a MiSeq Reagent Kit v3 (75 cycles) or NextSeq® 500 Sequencer with a High Output Kit v2 (75 cycles).

### Adapter Preparation

Each double-stranded adapter with a sticky end comprised two complementary oligonucleotides. D7 adapters have 4 bp sticky ends complementary to DNA cleaved by I-SceI; D5 adapters have 2 bp sticky ends complementary to DNA cut by MmeI. Each adapter was tagged by a central eight bp unique index (i7 and i5). The 5’ end of each D7_a and D5_a adapter bound the Illumina flow cell, the sequence 3’ of the indices were bound by Illumina’s Sequencing Primers. The oligonucleotide sequences are:

*D7_a: 5’-CAAGCAGAAGACGGCATACGAGAT[i7]GTGACTGGAGTTCAGACGTGTGCTCTTCCGATCT**TTAT**-3’*
*D7_b: 5’-AGATCGGAAGAGCACACGTCTGAACTCCAGTCAC[rc-i7]ATCTCGTATGCCGTCTTCTGCTTG-3’*
*D5_a: 5’-AATGATACGGCGACCACCGAGATCTACAC[i5]ACACTCTTTCCCTACACGACGCTCTTCCGATCT**NN**-3’*
*D5_b: 5’-AGATCGGAAGAGCGTCGTGTAGGGAAAGAGTGT[rc-i5]GTGTAGATCTCGGTGGTCGCCGTATCATT-3’*

Over-hanging bases are denoted in bold. The unique adapter indices (i7 and i5) are: D701: ATTACTCG; D702: TCCGGAGA; D703: CGCTCATT; D704: GAGATTCC; D705: ATTCAGAA; D706: GAATTCGT; D707: CTGAAGCT; D708: TAATGCGC; D709: CGGCTATG; D710: TCCGCGAA; D711: TCTCGCGC; D712: AGCGATAG; D501: TATAGCCT; D502: ATAGAGGC; D503: CCTATCCT; D504: GGCTCTGA; D505: AGGCGAAG; D506: TAATCTTA; D507: CAGGACGT; and D508: GTACTGAC.

Pairs of oligos (200 ng/μl each) were annealed to form adapters by incubating at 85°C for 10 min in 1x T4 DNA ligase buffer (NEB) before cooling gradually by transferring the heated block to the bench before dilution in 1x T4 DNA ligase buffer to working concentrations.

### Spike-in experiment to test sensitivity and quantification of mutant detection

Cells from 32 REMI mutants with defined insertion sites were grown in separately on SM plates in association with *K. aerogenes*. After washing in KK2 buffer, cells of each genotype were resuspended to 1×10^7^ cells/ml, and mixed at varying frequencies to generate four pools of cells (see Supplementary Table 3) where the relative frequency of ‘1000’ was 1×10^8^ cells, ‘100’ = 1×10^7^ cells, ‘10’ = 1×10^6^ cells and ‘1’ = 1×10^5^ cells. Genomic DNA from each of the four pools was extracted, and prepped for REMI-seq as described above before sequencing on an Illumina MiSeq with a MiSeq Reagent Kit v3 (75 cycles).

### Analysis of reads

First, 28 bp of adapter sequence was trimmed from each read as the extracted fragments are only 47 bp long. Next, each read was checked for the presence of the vector sequence [GC]AT[CG]CGTTGGA using an R Shiny app (https://github.com/NicoleGruenheit/REMI-seq-screen). If the sequence was detected, the script extracted the 6 bp index sequence, which is located 13 bp after the GATC or CATG, as well as the genomic tag including the DpnII or NlaIII site resulting in a 19/20 bp tag sequence. Then, each tag and index combination was counted and finally compared to a pre-computed lookup table. The orientation of the insert was determined using a comparison of the extracted tag to the upstream and downstream tag of each possible site together with the information whether the extracted index sequence was the left or the right side of the insert. The script also supplies basic statistics like the total number of reads per file, total number and percentage of reads that contain a tag, number of unique tags, number of tags in the inverted repeat (Chromosome DDB0232429, 2263132 to 3015703 is repeated between bases 3016083 and 3768654), and number of not unique tags. Then, raw read counts were normalised to make read counts comparable across samples using the total number of reads per sample and the total number of reads per insertion point (for upstream and downstream tag) was determined. Finally, insertion points were filtered for PCR artefacts (low occurrence of different index), and not unique tags (tags that cannot be uniquely assigned to one position are removed).

### Cell motility assay

Cells were re-plated from their original GWDI-bank growth 96-well plates into a new grid at the lower density of 5 ×10^4^ cells per well. Once the cell had adhered to the plate surface, they were imaged with on a CellInsight CX7 High-Content Screening (HCS) system and analysed using the cell motility package within the ThermoFischer HCS Studio Cell Analysis Software. For each strain, between 100-1000 individual cell images were captured at 2 min intervals for a minimum of 44 mins. The distance travelled between each time point was calculated for each cell and then averaged (mean) for the whole population. Each time value was then summed to give the Total Path Length for each strain. The measurements of two independent strains was averaged (mean) and compared to wild type. Significance differences were analysed using a T-test.

### Generation of mutant pool

Total number of transformants was estimated by counting the number of wells without any colonies in a subset of the 96 well plates. Cells were collected from the 96 wells plates after ^~^3-4 weeks incubation at 22°C, centrifuged for 2 minutes at 540 g at 4°C and resuspended in HL-5 medium including glucose and PVS. Cells were transferred to 14 cm petri dishes and incubated at 22°C. After 1 hour any dead cells were removed by changing the media. Cells were grown to confluency in HL-5 medium, resuspended at a density of ^~^5×10^7^ cells/mL in freezing media (50% Fetal Bovine Serum, 42.5% HL-5, 7.5% DMSO) and aliquots stored at − 80°C. For growth, cells were thawed directly into filter-sterilised HL-5 media, allowing cells to recover for 24 hours.

### Selection for growth in axenic culture

Cells from the mutant library (23,360 mutants) were seeded at a density of 2×10^5^ cells/ml on 10cm tissue culture plates and allowed to grow around 36 hours at 22°C (^~^4 generations) to reach confluency (3-4×10^6^ cells/ml) before being reseeded at 2×10^5^ cells/ml and grown for another 36 hours. Cells were frozen down for every 72 hours of continuous growth (^~^8 generations). Cells from a minimum of 5 plates were pooled and reseeded for each of the 2 biological replicates to mitigate bottlenecking of the mutant library (minimum 1×10^6^ cells reseeded = each mutant represented >40x). Each replicate was grown for a total of 80 generations. Genomic DNA was prepared from the mutant library following 24, 48 and 80 generations of axenic growth and processed for sequencing as described above.

### Selection for growth on *K. aerogenes*

The mutant library was initially thawed directly into filter sterilised HL-5 media and allowed to recover for 24 hours. Cells were then collected and washed twice in KK2 before being resuspended at 1×10^7^ cells/ml. 25μl of the suspension (2.5×10^5^) was then mixed with 400μl of an overnight culture of Ka and plated onto an SM plate. For each biological replicate of the screen, 4 plates were prepared in this way to prevent bottlenecking of the mutant library during serial transfer (1×10^6^ total cells = each mutant represented >40x). Following 48 hours of growth (approximately 10 generations of growth), when the cells had eaten most of the bacteria but had not yet begun to aggregate and enter development, cells from all 4 plates were pooled, harvested and washed by repeated centrifugation at 500g in KK2 until all the bacteria had been removed. Cells were then resuspended at 1×10^7^ cells/ml, and 2.5×10^5^ cells were replated serially on fresh bacteria to begin another ‘round’ of selection. An aliquot of cells were also resuspended at 5×10^7^ cells/ml in freezing media and stored at − 80°C.

### Analysis of mutant pools before and after selection

The mutant library was prepared for Illumina sequencing in triplicate as described above. Following sequencing, mutants were binned according to their mean normalised starting read counts (bin 100 = <100 reads, bin 1,000 = 100-1,000 reads, bin 10,000 = >1,000 reads). Following sequencing of mutant pools after selection, logfold change values relative to starting read count were calculated for each insertion mutant (Supplementary file 9). To allow comparisons of mutants across time, we normalised these data to have a mean of 0 and a standard deviation of 1 for each of the 3 bins of mutants. Mutants with a *z*-score >1.5 (i.e. >1.5 standard deviations from the mean of that bin) were considered to have an advantage under that condition, and mutants with a z-score < −1 a disadvantage. Mutants with fewer than 100 starting read counts that dropped out were discounted from this analysis because the technical drop out rate for these mutants was very high (Figure 4B). Hierarchically clustered *z-*score data across time for each of the screens were visualised using the ggplot2 package in R (Wickham, 2016).

### Growth mutant validation

A selection of axenic (7 mutants) and bacterial (13 mutants) mutants showing growth phenotypes was validated by growth competition assays with GFP-labelled AX4. To perform the growth competitions, cells of both AX4-GFP and competitor mutants were initially grown in parallel in tissue culture. Cells were then harvested, washed twice in KK2 and resuspended to 1×10^7^ cells/ml. To start each competition, mutant clones as well as the parental AX4 control were mixed 50:50 with AX4-GFP, and 2.5×10^5^ cells were then plated on either a single SM plate in association with 400μl *K. aerogenes* (bacterial growth competitions) or in tissue culture with 10mL HL5 (axenic competitions) in duplicate for 2 technical replicates per competition. Cells were allowed to grow together in for approximately 20 (axenic) or 40 (bacterial) generations. The relative proportion of GFP-labelled to unlabelled cells was scored at the start as well as the end point of the competitions by flow cytometry (Attune NxT Flow cytometer). Competition data was normalised to 100% and 0% labelling controls as well as to starting frequency and wild-type labelled vs wild-type unlabelled controls. Data is expressed as a change in frequency from 0.5 (see Supplementary file 10) from at least 2 independent biological replicates.

### Fluid uptake assay

The fluid uptake assay was performed as described in (Williams and Kay, 2018a), adapted to a 24-well dish format. Briefly, 2.5 × 10^5^ axenically growing cells were plated in triplicate on a 24 well dish. After settling onto the dish for 1 hour, the HL5 media was aspirated and the cells incubated for 1 hour with 0.5mg/ml TRITC-dextran (average Mw = 155kDa, Sigma Aldritch). The entire plate was then submerged and emptied twice in cold KK2 to quickly wash the cells, before cells from each well were collected in 1mL ice cold KK2 + 5mM sodium azide to prevent continued exocytosis. Median fluorescence intensity was then measured by flow cytometry (Attune NxT Flow Cytometer). Values were adjusted for background by subtracting the unlabelled control, and then normalised to control parental AX4 strain uptake (see Supplementary file 10), with at least 3 independent biological replicates performed per strain.

### Transcriptomes and GO analyses

To generate transcriptome data for axenically growing cells, we performed RNA-seq (Illumina). 2×10^7^ mid-logarithmic cells were collected and resuspended in 1mL TRIzol Reagent (Life Technologies) over consecutive days for a total of 3 replicates and stored at − 80°C. After thawing, samples were mixed with 200μl chloroform, incubated for 2 mins at room temperature and spun for 5mins at 16,000rpm at 4°C. The colourless aqueous phase was separated and RNA precipitated in 500μl isopropanol, mixed and spun 10mins at 16,000rpm at 4°C. The pellet was then washed in 1mL 70% ethanol, and finally dried and resuspended in 40μl RNase free ddH_2_O. RNA integrity was checked on an Agilent 2200 TapeStation. mRNA was then extracted by poly-A enrichment. RNA libraries were prepared using the Illumina TruSeq kit and sequenced using 100bp paired-end reads (150bp insert) on an Illumina Hiseq 4000. Reads were initially trimmed to remove remnants of Illumina TruSeq adapters and low quality basepairs using the trimmomatic package (Bolger, Lohse and Usadel, 2014). After this, any reads of less than 20 bases were discarded, as were any in which the average quality of the sequence was less than a phred score of 15. In all samples, more than 90% of the reads were retained. Trimmed sequences were then mapped to the AX4 *D. discoideum* reference genome using Bowtie2 (Langmead and Salzberg, 2012). A 750kb duplication on chromosome 2 of AX4 was masked to ensure that reads mapping to the genes in this region were not filtered out subsequently because they do not map uniquely to the genome. Sensitive end-to-end mapping parameters were used requesting the 10 best matches. Finally, any reads that mapped multiple times to the genome were excluded, using the mapping quality flag in the resulting SAM files. Mapped reads were sorted using samtools (Li, 2011) and converted to BAMfiles before reads mapping to annotated genes were counted using the RPKM_count.py script from the RSeQC package (Wang, Wang and Li, 2012).

To obtain bacterial growth transcriptome data, we downloaded the raw reads of RNA-seq data from the NCBI SRA database (accession numbers SRX271991 - SRX271998) for bacterially grown cells that used the same strain (AX4) under the same growth conditions as we used during the screen (clearing plates on SM agar in association with *K. aerogenes*) (Parikh *et al.*, 2010). To ensure that differences in gene expression are not due to different methods to map and count the reads, we subjected the reads to the same pipeline used for the samples grown in HL5 (see above). We then filtered out low read count genes (<100 mean read counts in both axenic and *K. aerogenes* conditions) and calculated an axenic expression index by dividing the mean axenic read count by the sum of the mean read counts of both conditions (1 = expression exclusively on bacteria, 0 = expression exclusively axenic) to compare the expression level of genes in different conditions with the growth phenotype. To generate gene lists of differentially expressed genes, we performed differential gene expression analysis using the R package DESeq2 (version 1.14.1)(Love, Huber and Anders, 2014). Genes were considered to be differentially expressed if they showed >2 fold difference in expression and an adjusted *p*-value <0.01 (Supplementary file 11).

To perform GO analyses on transcriptome and mutant lists, we used the GSEAbase R package (Morgan, Falcon and Gentleman, 2018) using a cutoff of *p*=0.05 for significantly over-or underrepresented GO terms. Gene lists for genes with phenotypes on *K. aerogenes* or HL5 were compared against a gene universe of genes from every mutant in the starting library. For differentially expressed gene lists the gene universe comprised all annotated genes in the *D. discoideum* genome (Eichinger *et al.*, 2005) (Supplementary file 12)

### Availability of Data

Raw reads of all samples were uploaded to the NCBI SRA database (PRJNA524784 and PRJNA524539). The R Shiny apps to analyse pools of REMI-seq mutants or grids can be found here https://github.com/NicoleGruenheit/REMI-seq-screen and here: https://github.com/NicoleGruenheit/grid_analyser. Detailed protocols, computational pipelines and further information can also be found at www.remi-seq.org.

---

**Supplementary Figure 1.**
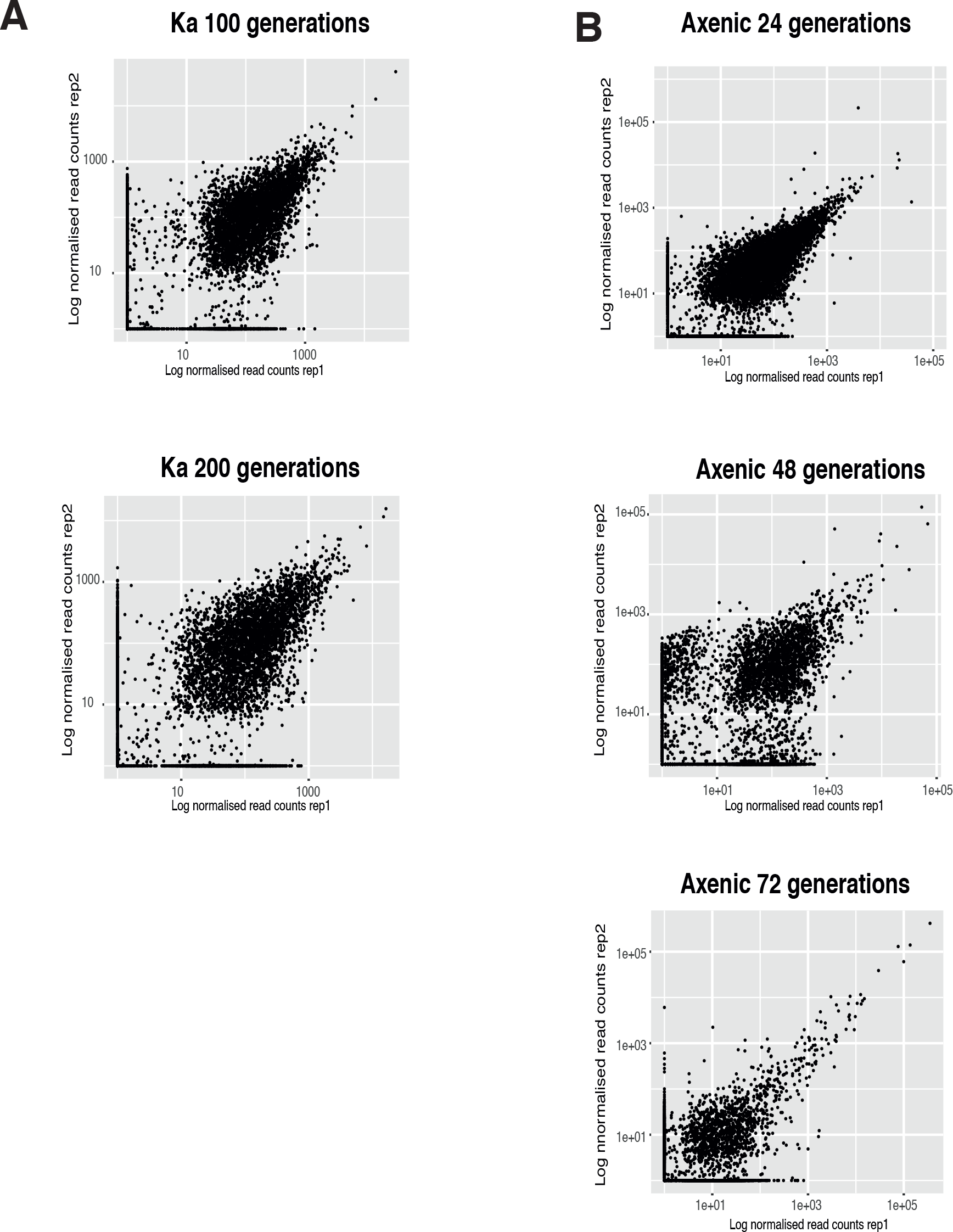
Read counts associated with each mutant are highly correlated between biological replicates from each round of selection after growth on bacteria (A) or in axenic medium (B).

**Supplementary Figure 2.**
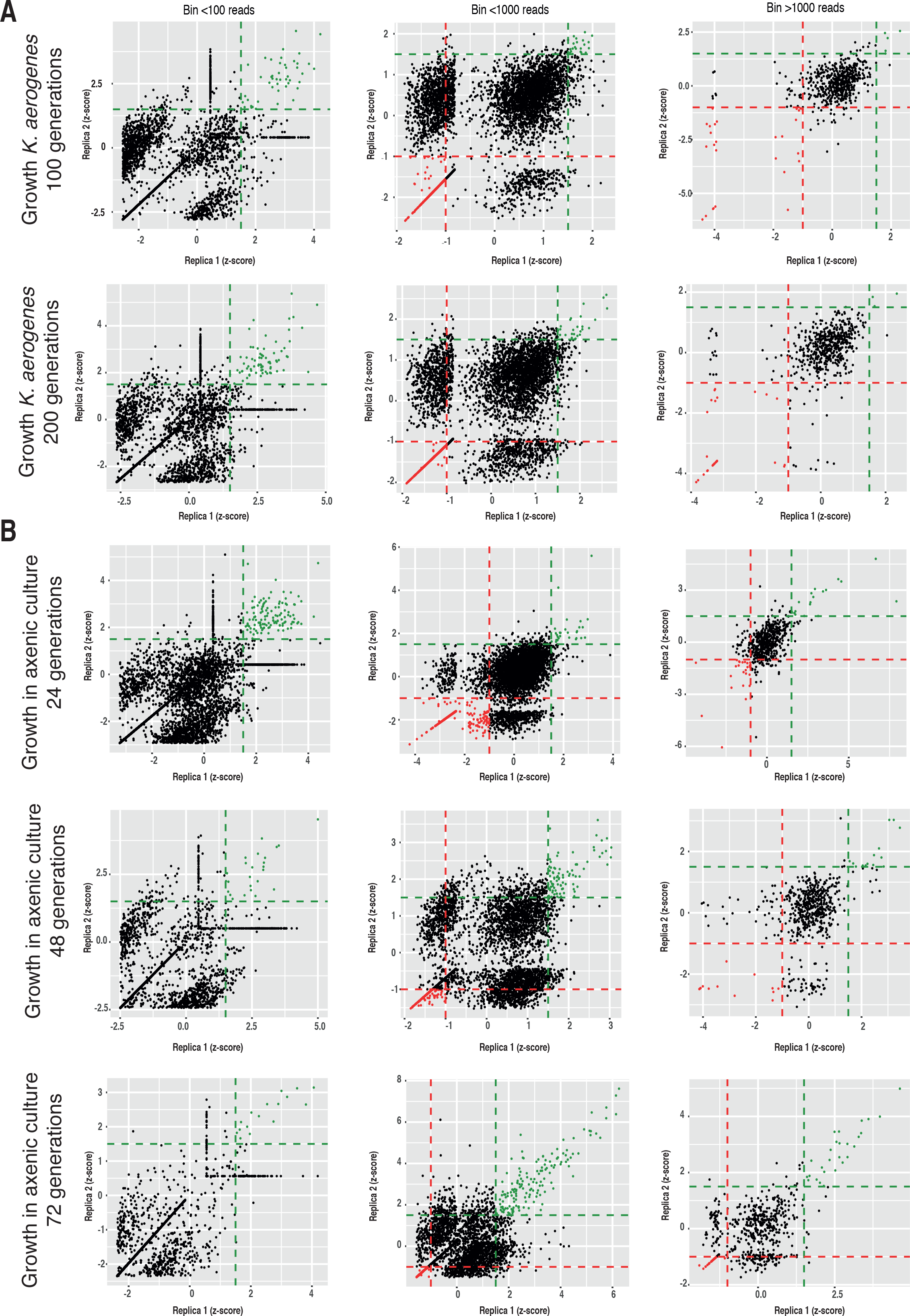
Identification of significantly enriched or depleted mutants from each round of selection after growth on bacteria (A) or in axenic medium (B).

**Supplementary Figure 3.**
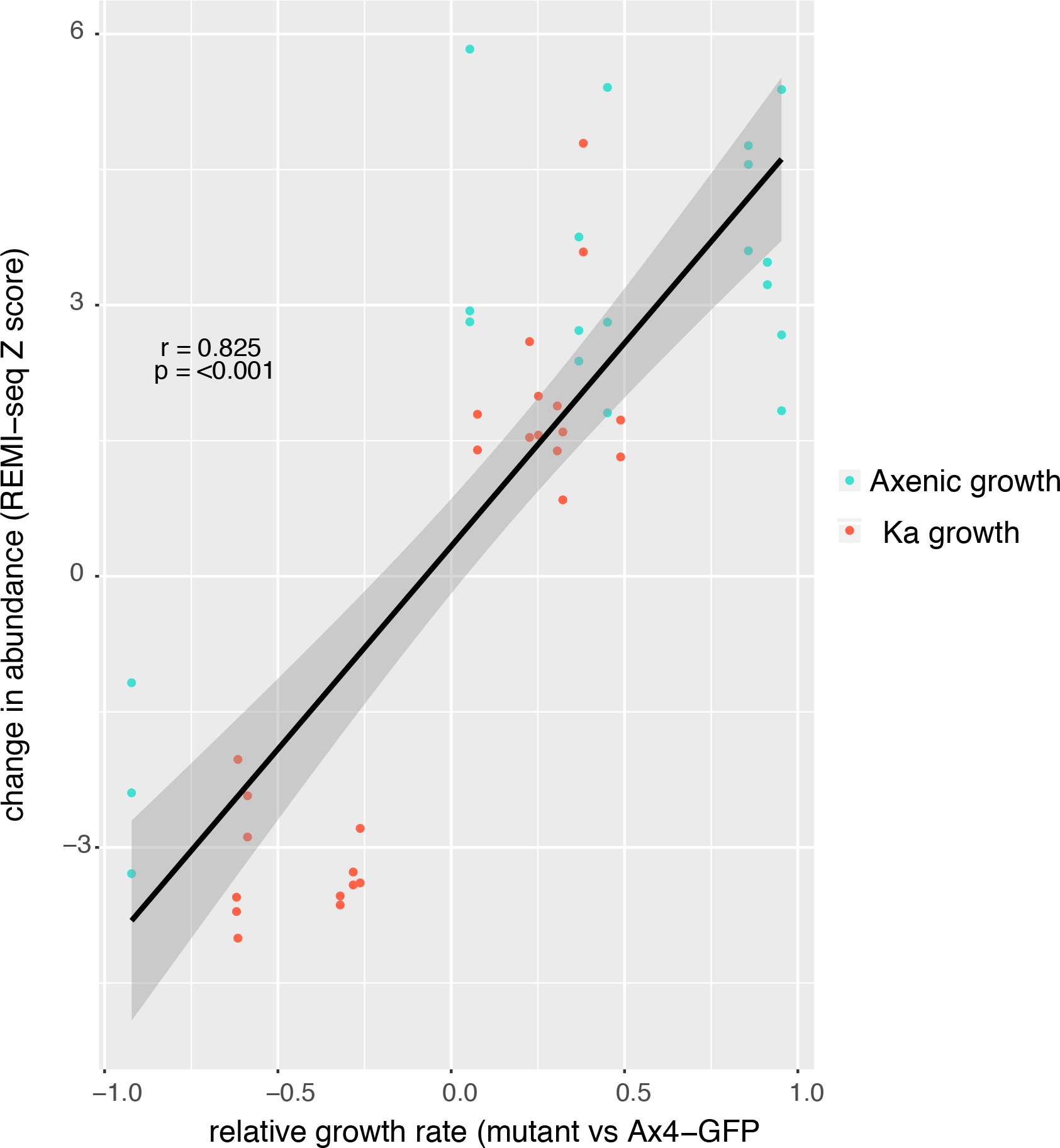
Correlation between the abundance of each mutant inferred from sequencing reads and the relative fitness (growth rate) of validated mutants across rounds of selection.

**Supplementary File 1: Sequence of the p-GWDI-G plasmid in SnapGene viewer format.** The sequence of the complete p-GWDI-G plasmid (4,488 bp) was annotated with the primers used for amplification of the insert as well as the left and right arms and the blasticidin resistance gene (bsr).

**Supplementary File 2: Sequence of the p-GWDI insert.** The sequence of the p-GWDI insert (1,593 bp) was amplified from the p-GWDI plasmids using the primers X and Y. It is centered around the blasticidin cassette. I-SceI and MmeI sites were integrated into the ends of this insert to enable the identification of the insertion site via next generation sequencing.

**Supplementary File 3: Lookup table for all possible insertion sites in the Dictyostelium genome.** Using the complete genome sequences of *Dictyostelium discoideum* the positions of all DpnII (GATC) and NlaIII (CATG) sites were identified as well as 20 bp up- and downstream of the recognition sites (genomic tags; Tab 1). Then, the number of occurrences of each tag in the genome as well as the location within or between genes was determined using this lookup table and the annotation file obtained from dictyBase. On top of this, the number of possible insertion sites was determined for each annotated gene (13,411; Tab 2) and all promoter regions (1,000 bp before the start codon). A short summary can be found in Tab 3.

**Supplementary file 4: Indices used for different p-GWDI plasmids.** p-GWDI plasmids are derived from the pLPBLP plasmid. The ends of both arms contain indices specific for each arm and plasmid. p-GWDI G1, G2, G3 and G5 have GATC sticky ends and p-GWDI C4, C6, C7, C8 have CATG sticky ends.

**Supplementary File 5: Table of individual mutants present in the pREMI grid.** Individual mutants were identified in 96-well plates using the REMI-seq technique (see methods). To identify the exact insertion points, the extracted tags were compared to the lookup table. The unique indices at the end of the inserts also allowed the determination of the vector version (p-GWDI G1 – G5; C6 – C8) as well as the orientation of the insert.

**Supplementary file 6: Mutants used in cell motility assays**

**Supplementary File 7: Table of individual mutants present in the pREMI pool.** Individual mutants were identified after sequencing a pool of approximately 20,000 mutants using the REMI-seq technique (see methods).

**Supplementary file 8: Table of insertion mutants used in spike-in study to test the quantitative range of REMI-seq**. 32 mutants with known insertion points were divided into four groups (A – D; Tab 1). Mutants were then mixed according to the experimental design table (Tab 2), where mutants were added to four different pools at varying quantities. Tags were then prepared using the REMI-seq protocol and sequenced on a MiSeq. Counts for each mutant (Tab 3) were normalised according to the total number of reads per sample.

**Supplementary File 9: Read count data before and after selections for growth in axenic culture or on bacteria.** Normalised read count data is shown for each detectable insertion mutants for each replicate of each condition, as well as log fold change values from the starting read count before the selections.

**Supplementary file 10: Growth competitions and fluid uptake assays.**

**Supplementary file 11: Differential gene expression analysis on axenic vs bacterial growth**

**Supplementary file 12: GO term analysis of genes associated with growth in axenic culture or on bacteria**

## Acknowledgements

This work was supported by a Wellcome Trust Biomedical Resources Grant to AJH and CT, and a Wellcome Trust Investigator Award to CT. We thank Angela Marchbank and Dr Nicholas A. Kent in the Cardiff School of Biosciences Genomic Research Hub, Dr Andrew Hayes in the Genomic Facility at the University of Manchester and Dr Tony Brooks in the UCL Genomics Facility for their expert advice.

