## Supplementary file 6 for "REMI-seq: Development of methods and resources for functional genomics in *Dictyostelium*"

| **Gene name** | **Putative Function** |
| --- | --- |
| kxcB | serine/threonine protein kinase with RhoGEF, DH and PH domains |
| docA | DOCK180 protein homologue |
| zizA | zizimin- related DOCK family protein member |
| vilC | villin-like protein C |
| elmoE | ElmoE forms a complex with DocC and ZizA |
| rapgap1 | **Rap**GTPase, **G**uanosine triphosphatase-**A**ctivating **P**rotein 1 |
| Roco10 | LRRK family protein kinase |
| DDB_G0277675 | RasGTPase-activating protein |
| DDB_G0283827 | WASP-related protein C |
| DDB_G0289829 | Unknown |
| DDB_G0277997 | Neuroblastoma-amplified sequence |
| DDB_G0277165 | CAMKL homologue, BRSK protein kinase subfamily |
